# Transmission of human-associated microbiota along family and social networks

**DOI:** 10.1101/540252

**Authors:** Ilana L. Brito, Thomas Gurry, Shijie Zhao, Katherine Huang, Sarah K. Young, Terrence P. Shea, Waisea Naisilisili, Aaron P. Jenkins, Stacy D. Jupiter, Dirk Gevers, Eric J. Alm

## Abstract

The human microbiome, described as an accessory organ because of the crucial functions it provides, is composed of species that are uniquely found in humans^1,2^. Yet, surprisingly little is known about the impact of routine interpersonal contacts in shaping microbiome composition. In a relatively ‘closed’ cohort of 287 people from the Fiji Islands, where common barriers to bacterial transmission are absent, we examine putative bacterial transmission in individuals’ gut and oral microbiomes using strain-level data from both core SNPs and flexible genomic regions. We find a weak signal of transmission, defined by the inferred sharing of genotypes, across many organisms that, in aggregate, reveals strong transmission patterns, most notably within households and between spouses. We find that women harbor strains more closely related to those harbored by their familial and social contacts than men; and that transmission patterns of oral- and gut-associated microbiota need not be the same. Using strain-level data alone, we are able to confidently predict a subset of spouses, highlighting the role of shared susceptibilities, behaviors or social interactions that distinguish specific links in the social network.

---

Host-specificity rather than generalist life histories dominate in the colonization of the gut^3^. Thus, colonization likely depends on direct interpersonal interactions where individuals are exposed to other humans’ microbiota. Nevertheless, the extent to which regular, repeated bacterial exposures result in transmission is unknown. Mother-to-child transmission can be detected early in life^4,5,6^, but these patterns fade, whereas other factors—environment^*7*^, behaviors and genetics^8^ impact the strain-level composition of each adult’s microbiome^9,10^. The human microbiome remains remarkably stable in composition over days^11^ and even years, at the level of strains^10,12^, raising the question: do we exchange oral and gut commensals with our closest family and friends?

Here, we take advantage of rich familial and social network data obtained as part of the Fiji Community Microbiome Project (FijiCOMP) (Figure 1a, Supplementary Tables 1-2) to explore the role of transmission in human populations with strain-level resolution. Our data consists of shotgun metagenomic sequences from 287 people living in 5 agrarian villages in the Fiji Islands (Supplementary Table 3, Supplementary Table 4). Paired gut and oral microbiome samples were deeply sequenced to enable molecular epidemiological analyses. The presence of locally endemic bacterial disease suggests that commensal bacteria may also spread widely within the community. Due to the relative isolation of these villages and the reliance on local food and water, we hypothesized that with comprehensive sampling of eligible individuals in each village, we could capture all human sources and sinks of human-associated bacteria, enabling the tracking of strains within this comparatively ‘closed’ network.

**Figure 1.**
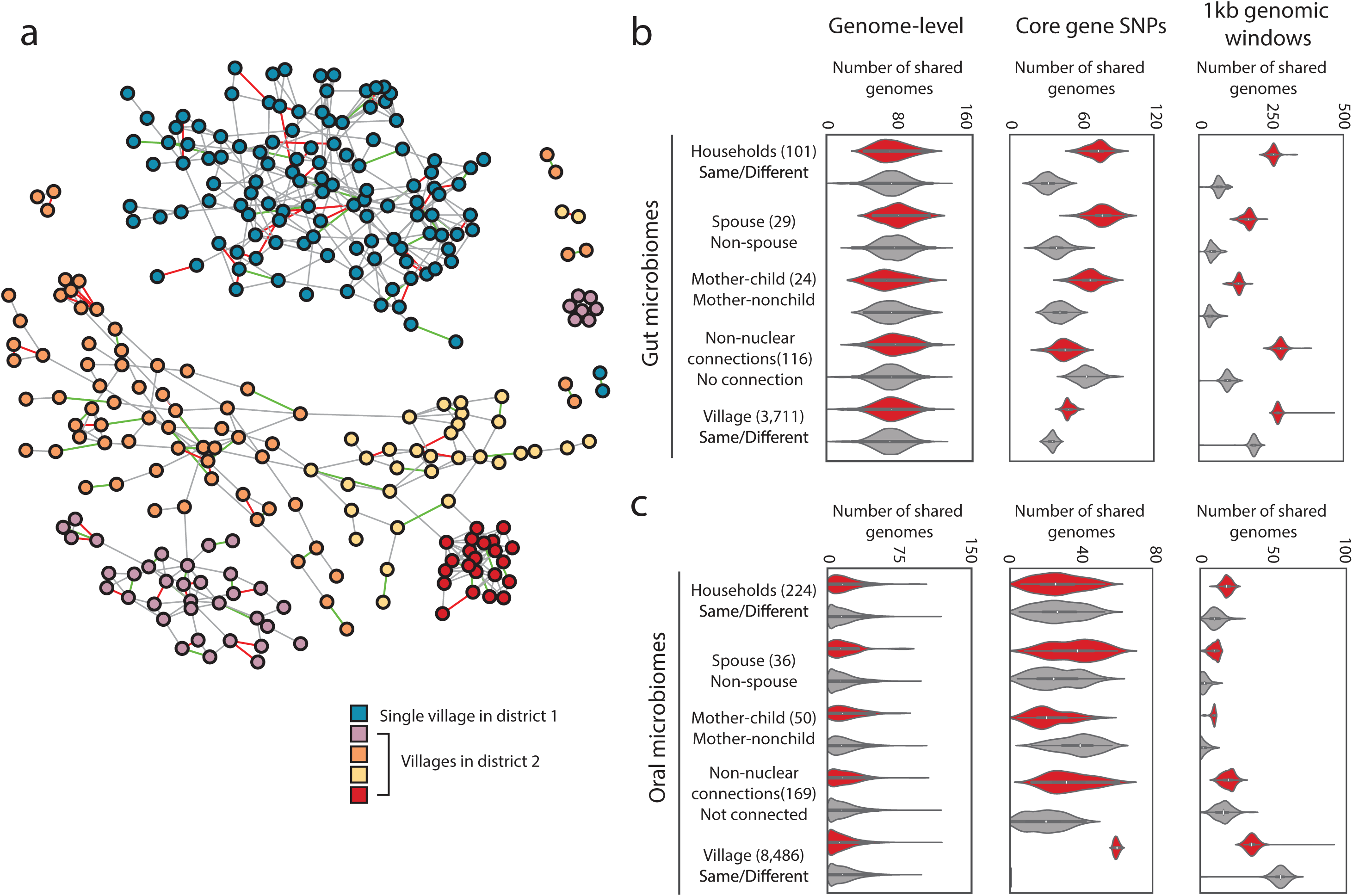
Household membership results in shared bacterial lineages. (a) The family and social network of the FijiCOMP cohort, colored by village membership. Four villages are in the same district whereas the fifth village is in a different district. Spousal relationships are designated by edges colored red, whereas mother-child relationships are designated by green edges. Gray edges represent all other familial or social network relations. (b,c) In the gut (b) and oral (c) microbiome samples, the number of shared genomes (left), the number of genomes with shared core gene SNP profiles, determined by Manhattan distances (middle), and the number of genomes sharing flexible genomic regions, determined by 1kb genomic windows (right) significantly associated with pairs of linked rather than unlinked members of the social network. A ‘genome’ refers to each assembled sets of core proteins for each species (left and middle), or to each assembled LSA partition (right). Any connection refers to friendship or distant familial connections in the network, excluding nuclear family and household connections. Full p-value distributions for the distributions shown are in Supplementary Figure 4. The violin plot distributions represent results from comparing the linked pairs in a given social network (red) or the shuffled network (gray) with *N=*100 independent sets of the unlinked pairs obtained by bootstrapping. Whiskers inside the violin extend to points within 1.5 interquartile ranges (IQRs) of the lower and upper quartile for a distribution, and center points represent its median. The numbers of linked pairs for each network (stool/saliva) are as follows: household (101/224); spouse (29/36); mother-child pairs (24/50); any connection (116/169); village (3,711/8,486).

The bacteria present in the FijiCOMP microbiomes are largely distinct from those in existing databases^13^ resulting in poor read alignments to reference genomes (Supplementary Figure 1). Therefore, we binned reads derived from oral or gut microbiomes using Latent Strain Analysis^14^, and *de novo* assembled a set of draft genomes (Supplementary Table 5). There were little to no detectable differences in species-level sharing than expected by chance across any relationship type in either the gut or oral microbiome samples (Figure 1b,c, Supplementary Figure 2), a finding at odds with that of households in Kenya^15^, Israelis^16^ or metropolitan Americans^9^, yet one that may reflect the high contact rates between individuals in this cohort.

To achieve strain-level resolution within individuals’ microbiomes, we employed two orthogonal approaches, focusing on either polymorphisms in core proteins or the presence/absence of flexible genomic regions. The former involved aligning sequencing reads to sets of core genes from each of the assemblies (Supplementary Table 5), similar to several established methods^6,7,17^, adjusted for use within the context of a social network. Specifically, we calculated the Manhattan distances between pairs of individual’s putative genotypes, inferred by the dominant SNP at each polymorphic position in the alignment. For individuals in the same village, household members or non-nuclear connections, we compared the distances for each genome of all connected pairs and a balanced random subset of unconnected pairs; whereas we simply shuffled the associations of spouses and mother-child pairs. We performed 100 bootstraps of the unconnected pairs or shuffles, each time tallying the number of genomes for which the median Manhattan distance was lower in connected individuals versus unconnected (Figure 1b,c). We next implemented an alternate strategy, largely based on the previous observation that flexible genomic regions may be highly personalized^18^. Coverage of one kilobase windows of contigs over 10kb were compared across pairs of individuals. Shared genotypes were defined by the complete lack of outlying 1kb regions present in one individual and absent in the other (Supplementary Figure 3). We tallied the number of assembled genomes more frequently shared in each relationship type in over 100 shuffles or bootstraps, again controlling for class imbalances, resulting in the distributions in Figure 1.

Transmission, loosely defined by shared inferred genotypes, has been observed for strains within the gut microbiomes of mother-child pairs^19^, albeit most notably in the first year of life^4,5,6^, in cases where fecal material was used for transplantation^17,20^, and between twin-pairs^10^. Within the village setting, we are unable to determine whether strain transfer is direct or indirect, or from a common source, nor can we infer its directionality. However, we refer to the presence of shared genotypes as ‘transmission’ as the putative explanation for the observed patterns. Here, consistent patterns of transmission were revealed across individuals’ social networks in both gut and oral microbiomes, independent of the metric used (Figure 1b,c, distributions of p-values in Supplementary Figure 4). Household members showed high levels of strain similarities in their gut microbiomes, across mother-child pairs and, most notably, among spouses, who share no genetic relatedness. The length of cohabitation was positively correlated, albeit weakly, with the measure of strain dissimilarities (Supplementary Figure 5), which may reflect long-term changes in intimacy or lifestyle.

The signal varies across our two metrics, potentially highlighting interactions in which organisms versus mobile genetic elements are transmitted between individuals. Using a set of gut microbiome mobile genes previously identified in the FijiCOMP cohort^13^, we find mobile genes weakly shared between spouses (Supplementary Figure 6). Using strain-level metrics, the transmission signals are robust. Transmission within villages in both gut and oral microbiomes was detectable in core gene SNPs even when we rarefied the number of village pairs from over one-thousand down to 10 pairs each of connected and unconnected individuals (Supplementary Figure 7). Furthermore, our results were consistent even when we reduced the number of genomes considered using only those LSA-informed assemblies with low putative contamination (Supplementary Figure 8). In all cases, shuffling network relations, while retaining network architecture, ablated observable transmission patterns (Supplementary Figure 9).

We next examined the contributions of specific organisms, as familial transmission has been previously observed for certain gut and oral commensals^8,21,22,23^. There was no consistent signal of transmission across any single phyla (Supplementary Figure 10). Instead, each pair of connected individuals had a unique signature of shared organisms (Figure 2a,b, Supplementary Figures 11-18), suggesting that transmission may be largely driven by chance events and indirect transfer. Interestingly, the fidelity of our LSA-informed assemblies did not strongly impact our results, as transmission may still be observed even if core genes are shuffled between assemblies (Supplementary Figure 19), supporting the notion that signatures of transmission are distributed broadly over many strains. Overall microbiome functional profiles also failed to capture transmission signals (Supplementary Figure 20), though this does not negate the potential contributions of individual virulence- or transmission-associated genes contributing to transmissibility. We hypothesized that perhaps the abundance of each organism would be indicative of its overall transmissibility, favoring a mass-action model of transmission, yet this was not the case (Figure 2c,d).

**Figure 2.**
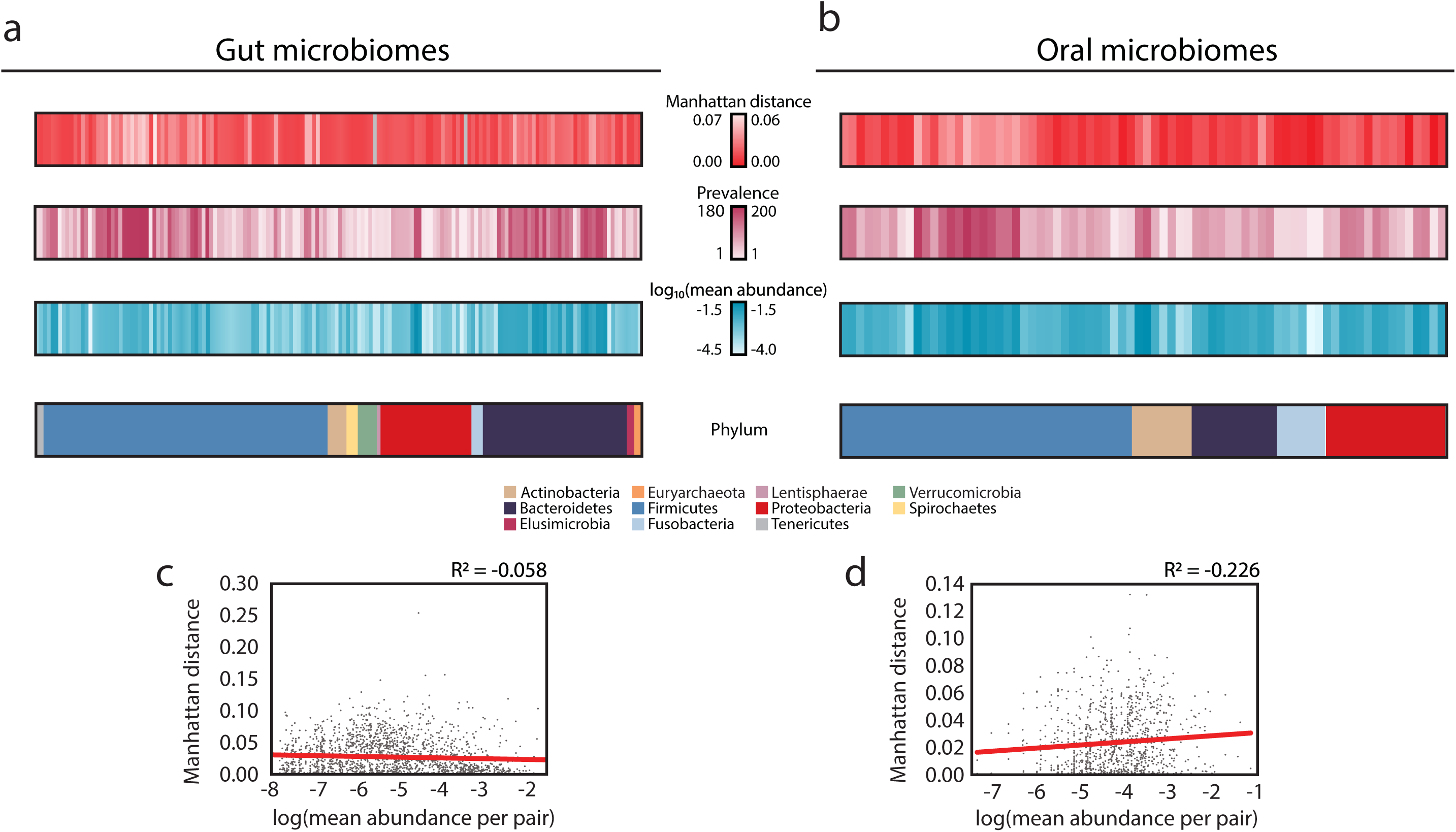
Organisms vary in their transmissibility across the social network. (a,b) The mean Manhattan distance, prevalence (number of individuals who harbor that organism), log_10_(mean abundance) and phyla are plotted for organisms in the (a) gut (*N=*1,988) and (b) oral microbiomes (*N=*1,111) of spouses. (c,d) The mean abundance of each organism across each pair of individuals is plotted against the Manhattan distance of that organism for that pair of individuals in the (c) gut and (d) oral microbiomes. Linear regressions are plotted in red.

These findings lead to an apparent paradox: if most bacteria are transmitted directly between members of the community, then why don’t we observe clearer patterns of transmission? We believe there are several factors that contribute to the ‘diffuse’ signal for transmission observed across this population. First, despite this relatively ‘closed’ network of individuals, there are inherent difficulties in capturing the full range of individuals’ contacts and exposures. Our best approximations of direct transfer may be far from actual events, where indirect transfer between individuals outside of the network or transmission from unknown and unsampled environmental reservoirs may play a consequential role. Second, we focus on a snapshot in time, not knowing *a priori* what types of interpersonal interactions result in transfer nor whether transmission occurs during particularly volatile points in an individuals’ microbiome history. Third, despite our achieved sequencing depth, perhaps longer-read sequencing or a massive increase in sequencing depth is required to achieve greater strain resolution. We reached the limit of detecting transmission when we rarefied samples to 5 million reads (Supplementary Figure 21). Finally, this community may actually be more prone to transmission with a wide range of community members, even when compared to other non-industrialized populations. This is best illustrated by regular gatherings to drink kava, where a communal vessel and cup are shared.

Borrowing from the framework of disease ecology, we sought to test the impact of specific individuals within the social network on overall network-level transmission. ‘Superspreading’ is a phenomenon observed for the transmission of diseases such as severe acute respiratory syndrome (SARS) and human immune-deficiency virus (HIV), where the majority of the transmission observed is attributable to a relatively small number of people^24^. Across our cohort, there were detectable differences in transmission per individual of both stool and saliva (Figure 3a-c,e, Supplementary Figure 22). Since we cannot determine the direction of transmission, we refer to this phenomenon as ‘supersharing’ in this cohort. Supersharing was largely agnostic to the individuals’ read depth, once a threshold is achieved for obtaining accuracy in Manhattan distances (Supplementary Figure 23). Interestingly, individuals who were strong supersharers of gut microbiota were not the same as those of oral microbiota (Figure 3g), revealing differences between the transmission routes of commensals. There was also no specific association with individuals’ overall sharing and their network positions, either in terms of the number of connections (‘degree’) or the centrality (measured by ‘betweenness’) (Supplementary Figure 24).

**Figure 3.**
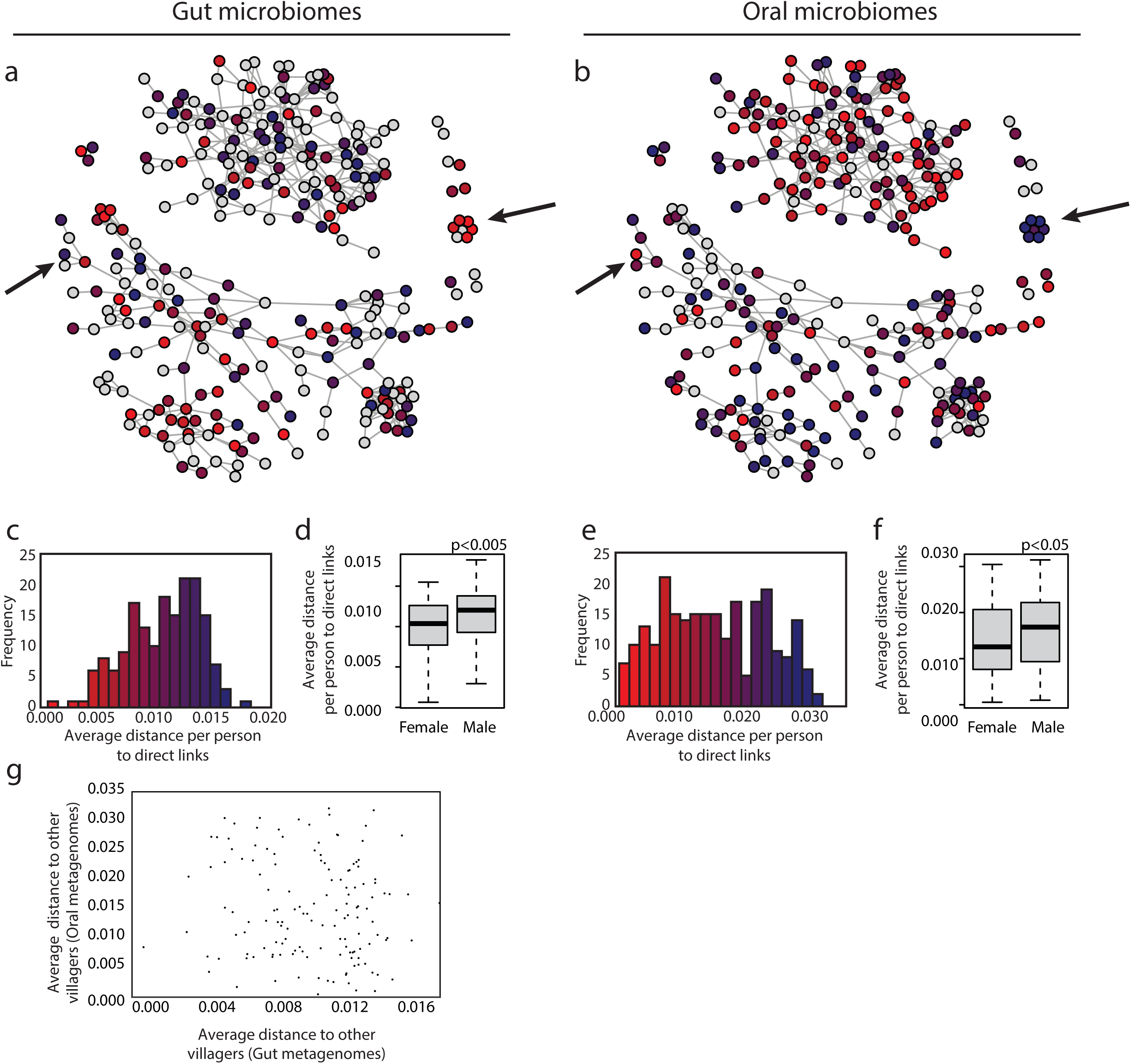
Some individuals are ‘supersharers’. (a,b) For each person in the network, the average distance, defined as the median of mean Manhattan distances across all genomes to all directly connected individuals, is plotted for organisms within the (a) gut and (b) oral microbiomes. Arrows point out examples in which individuals’ sharing patterns are different for gut and oral microbiota. The red and blue in plots (a) and (b) match the values plotted in parts (c) and (e), respectively. (c,e) The distribution of average distances (median of mean Manhattan distances) for each individual to all of their directly connected individuals is plotted for female and male individuals’ (c) gut (*N=*173) and (e) oral microbiomes (*N=*243). (d,f) A histogram of the average distances (median of mean Manhattan distances) for each individual to all of their directly connected individuals is plotted for individuals’ (d) gut (*N=*173) and (f) (*N=*243) oral microbiomes. Boxes indicate the upper and lower quartiles, whiskers extend to highest and lowest values excluding outliers, and center lines indicate medians. P-values were obtained from one-tailed Wilcoxon rank sum tests. (g) Each individual’s median of mean Manhattan distances to all individuals within the same village is plotted for their gut and oral microbiomes (*N*=142).

Surprisingly, sharing of both gut and oral microbiota was more associated with females in the network (p<0.005 for gut microbiomes, p<0.05 for oral microbiomes, Pearson correlation, Figure 3d,f, Supplementary Figure 25), yet had no relationship with age (Supplementary Figure 26). Although gender-related differences in pathogenic bacterial transmission are well known, as are the myriad factors that affect exposure and susceptibility^25^, these are less well understood for commensal microbiota with no clear mechanisms of transmission. Nevertheless, exposure risks may be associated with occupations and behaviors that are highly gendered within this cohort (housekeeping, p<10^−15^; farming and fishing, p<10^−15^, caring for ill family members, p<0.05; and soap usage p<0.05, chi-squared test). It remains to be determined how the transmission observed in this low-income, agrarian population would compare to a population living in an industrialized nation, where interventions such as the use of antiseptics, disinfectants and antibiotics, sanitation infrastructure and food safety restrictions, may influence the transmission of commensal bacteria.

We next asked whether strain-level information alone could be used to predict specific social relationships. We implemented a machine learning approach that utilized organism abundances, core SNP profiles, flexible region similarity or combinations thereof, without considering demographics. Our household predictions were moderately accurate (AUC = 0.64±0.02 and 0.61±0.01, for gut and oral microbiomes, respectively), whereas our model to predict spousal relationships performed better (AUC = 0.70±0.03 and 0.72±0.02, for gut and oral microbiomes, respectively) (Figure 4, Supplementary Figure 27). Despite the poorer overall performance of our household models, the predictions appeared dependent on the network structure, as all of the relationships within some households were accurately predicted, in both gut and oral samples. Remarkably, our model reveals that close to 25% of spouses are exceedingly easy to predict with high confidence (Figure 4c,d). Within the household network, some of these spousal pairs were obscured, highlighting the subtle nature of these transmission signals. Why certain couples are easier to predict than others is unknown, but may reflect shared susceptibilities, specific behaviors, or the relative importance of extra-marital relationships. Interestingly, spouses have been found to share immune repertoires^26^ and households display family-specific signatures^27^, providing evidence for shared exposures.

**Figure 4.**
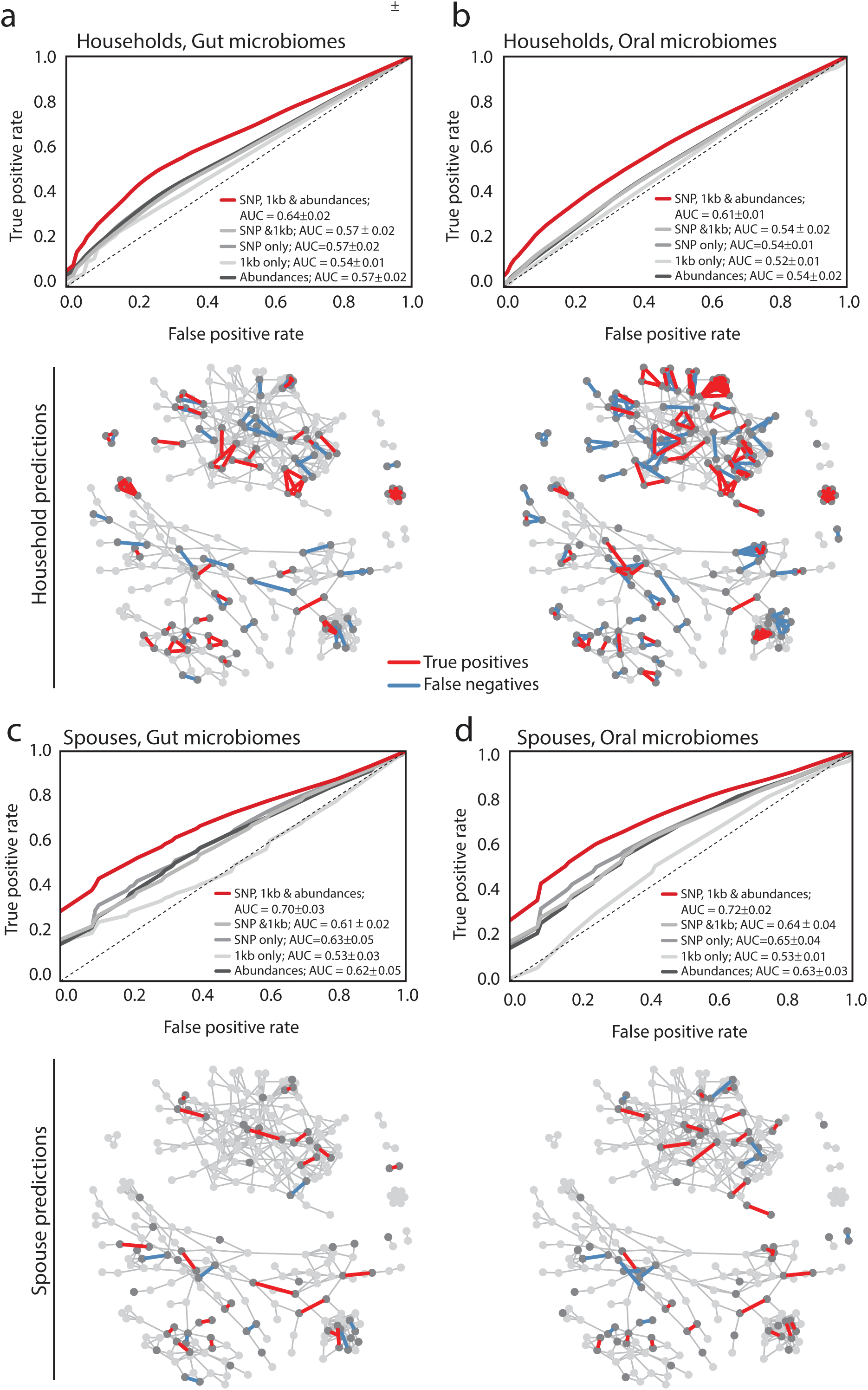
Machine learning predicts a subset of spouses with high confidence. (a,b) ROC curves for a random forest model predicting household membership based on shared (a) gut or (b) oral microbiome strain-level data are plotted for models using SNP profiles, shared flexible regions, both, or both with organismal abundances. Random forest models were constructed from 1,000 decision trees and without constraint on maximum tree depth. The dotted line shows an ROC where false positives equal false negatives. The legend reports means and standard deviations for each classifier’s Area Under the Curve (AUC). (c,d) The social network plotted with predicted true positive household pairs and false negative household pairs using gut (c) or oral (d) microbiome data. Arrows point to examples of either families in which everyone in a household can be confidently predicted. (e,f) ROC curves for a random forest model predicting household membership based on shared (e) gut or (f) oral microbiome strain-level data are plotted for models using SNP profiles, shared flexible regions, both, or both with organismal abundances. Random forest models were constructed from 1,000 decision trees and without constraint on maximum tree depth. The legend reports means and standard deviations for each classifier’s AUC. (g,h) The social network plotted with predicted true positive household pairs and false negative household pairs using gut (g) or oral (h) microbiome data. Arrows point to examples of either families in which everyone in a household can be confidently predicted.

Although it is well-established that shared environments significantly affect the gut microbiome composition and phenotype of isogenic mice^28,29^ and that social interactions shape wild primate microbiomes^30^, this work opens the door to understanding the process of transmission and its implications in human society. Within this small community of individuals with relatively homogeneous living environments, diets and microbiomes, bacterial DNA alone can be used to accurately predict certain intimately linked pairs of individuals. Our research begins to tease apart relevant transmission patterns evident in a social network and a role for gender in commensal transmission, revealing that long-term intimate interactions that occur later in life, such as through marriage or co-habitation, can result in stochastic transmission events in both the gut and oral microbiomes. Given the wide array of microbiome-associated health conditions, this study further hints at the possibility that diffuse transmission patterns of pathogenic or protective commensals may contribute to individuals’ overall health status.

## Methods

### Social network construction

The Fiji Community Microbiome Project (FijiCOMP) consisted of interviewing and sampling the gut and oral microbiomes of almost 300 individuals living in 5 village communities in two districts approximately 50 miles away from one another on Vanua Levu in the Fiji Islands. The sampling all took place within a 4-week period, each village taking approximately 1-2 weeks. IRB approval was received from Institutional Review Boards at Columbia University, the Massachusetts Institute of Technology and the Broad Institute and ethics approvals were received from the Research Ethics Review Committees at the Fiji National University and the Ministry of Health in the Fiji Islands. Informed consent was obtained from all study participants.

As part of the survey, each head of household was asked to draw their family trees, including all members of their household, even if they are not related. Individuals were specifically asked to name their spouse, if married, and the number and ages of their children. We inferred the number of years a married couple lived together by the age of their oldest child. We excluded six of the 63 couples from our analysis of the time they lived together because either they did not have any children or their children’s ages were inconsistent (for example, if children came from a previous marriage). As houses commonly have names rather than specific addresses in these villages, individuals were asked the name of the house in which they live. Individuals’ responses were cross-referenced for consistency and ambiguous links were removed from our analysis. Minor discrepancies, such as slight differences between spouses in the reporting of their children’s ages that differed by 1 year, were permitted. Individuals were further asked to provide the names of 5 individuals with whom they spent the most amount of time. Although the individuals mentioned the type of relationship (*e.g.* mother/child, cousin, sister-in-law, friend, classmate, churchmate *etc.*), these relationship types were not solely relied upon to define a particular relationship type. In a small number of examples, individuals cited social interaction with a third party whose identity could not be verified, and were therefore not included in our analysis. Additionally, some individuals mentioned siblings or parent/child relationships that could not be verified, so these were also counted as merely social interactions. This resulted in 489 unique social/familial interactions, in addition to household-level interactions. For the purposes of anonymity, individuals’ ages were rounded to the nearest 5-year increment and the number of children per person was not reported. Not all children of each family were surveyed, either because the children did not meet the inclusion criteria (they needed to be at least 8 years of age) or because they were inaccessible during the time when we were sampling. Social network was plotted using R package igraph (v.1.0.1). Network metrics (i.e. betweenness, degree) were calculated using igraph standard functions.

Additional information was obtained from all participants including having individuals name their occupation (of which domestic duties, farmer, and fisherman were all possible answers), whether the individual had cared for a sick family member in the past year, and whether they used soap (with possible answers: always, sometimes and never).

### Alignments and identification of single nucleotide polymorphisms

We calculated the Manhattan distances between the dominant SNPs within pairs of individuals’ core gene alignments. This involved aligning each individuals’ reads to core genes in the assembled LSA partitions, extracting polymorphic loci, and determining the dominant allele at each locus. For each pair of individuals, we computed the Manhattan distance at each locus, averaged this distance across loci, and computed this quantity for every partition/genome. These distances were then used for the network comparisons described in the ‘Network comparisons’ section.

More precisely, quality-filtered, dereplicated metagenomic datasets (on average, over 52 million and 10 million reads for our gut and oral microbiomes, respectively), devoid of human genetic material (filtered as described in Brito et al., 2016^13^), were partitioned prior to assembly using Latent Strain Analysis^14^ according to covarying kmer content across samples. Read partitions were then assembled using Velvet^31^. Sets of core genes were identified using AMPHORA2^32^. Core genes were assigned taxonomies using genera-level best hits using BLAST+ against the NCBI NR database. Partitions with complete (31 single-copy genes for bacteria) or near-complete gene sets of AMPHORA genes deriving from the same genera were retained for analysis (Supplementary Tables). If a core gene set contained more than 2 of the same assembled gene, we removed both copies of that gene.

Each individuals’ samples were then aligned to the sets of core genes using BWA-MEM^33^. Reads were subjected to more stringent trimming using TRIMMOMATIC^34^ (in addition to trailing low quality bp (quality < 4), we also implemented a sliding window, trimming when the quality < 15). Reads were then aligned to regions that included 1 read-length (100bp) downstream and upstream of each core gene to avoid edge effects within the alignments. 100bp from each end of the alignment, regardless of whether the gene was positioned at the end of the contig, was then trimmed from the final pile-up. Reads were filtered to retain those with greater than 40% of the length aligning at 90, 95, 97 or 99 percent identity. A lower cut-off was chosen to capture a wide variety of strains for each alignment. Setting a lower threshold would be more inclusive of strains more distantly related to the reference, which would only obfuscate a signal for a given species should it include too distant strains. Previous work^20^ estimated the species boundary at approximately 85-90% identity in core genes (analogous to ∼97% in the 16S rRNA gene). 90% identity also resulted in the most consistent coverage across core genes, and it was therefore chosen for all subsequent SNP-level calculations. Reads with soft- or hard-clipping were removed. To further validate our gene sets, we filtered out genes with abnormal coverage relative to the rest of the gene set. We expected the depth of each gene to be uniform across a genome, and sequencing depth to be Poisson distributed at each locus. To avoid including genes within a species’ genomes that recruiting abnormal numbers of reads compared to the remainder of the genome (and thus more likely to be recruiting reads from other species), we computed a chi-squared goodness of fit test for each gene between the empirical coverage distribution and the equivalent Poisson distribution of the same mean. Genes with median p-value lower than 0.05 across subjects were discarded from any subsequent analysis. Results were mostly bimodal, where most genes fit the equivalent Poisson distribution very well, giving us confidence that reads were being recruited uniformly across the full length of the considered genes.

To calculate genome-wide statistics (Figure 1, left), we built a table of the median coverage across the SNP tables within the core genes, across different assemblies. Then, for each pair of people, we counted the number of these genomes that they shared, and compared that between related and a balanced set of unlinked pairs.

Polymorphic loci were then identified from the alignment, resulting in a counts matrix for each genome containing read counts for each allele at each locus in each individual. We retained the dominant allele for each individual (the allele with the highest number of read counts) at each site, then then computed the Manhattan distance between each individual’s dominant allele at each site, and averaged these distances across each genome to obtain an average Manhattan distance per SNP for each genome in a given pair of individuals. For each pair of individuals in a given social network (e.g. same household), this average Manhattan distance per SNP was computed for every genome, and the median distance for a given genome compared to the median distance observed in unrelated pairs of individuals. This calculation is described in more detail in the ‘Network comparisons’ section.

As a comparison, we also ran the quality-filtered forward metagenomic reads through the MetaPhlan2^36^ pipeline.

### Abundance comparisons of 1kb windows in assembled genomes

Contigs under 10kb were removed from LSA-assembled draft genomes. Reads were aligned to contigs with 95% identity. Reads with hard and soft clipping were removed, as were Supplementary alignments. We only considered pairs where both individuals had a median coverage of 10 or more across the genome. 1kB regions were considered present in an individual and absent in another if its coverage was greater than the median in the first individual; and lower than one thousandth of the median in the other. Pairs of individuals were considered to share the same strain if there were no such 1kB regions across the entire genome (i.e. all regions were either present or absent in both individuals) and that it was present with a median coverage of 10 or more in both individuals.

### Mobile genetic element analysis

For Supplementary Figure 6, we used the abundances of mobile genes identified in Brito *et al.* 2016 to determine whether there was a transmission signal. We calculated the Jensen-Shannon divergence between all pairs and compared the number of pairs within each group with a balanced, subsampled group.

### Functional contribution to transmission

Genes in the LSA-assembled genomes were first clustered at 90% identity using CD-HIT^37^. Representative genes were then annotated using DIAMOND^38^ against the Kyoto Encyclopedia of Genes and Genomes (KEGG) database (release 73.0). Abundances for each gene were then calculated as the median read depth across genes with over 85% coverage. Abundances were summed for each functional gene family (represented by a single KO number). For each pair, the Jensen Shannon divergence was calculated.

### Network comparisons

Network comparisons on the mean pairwise SNP distance were performed by comparing the median value of the mean pairwise distance per SNP in related pairs with those in unrelated pairs for each genome. If a genome’s median pairwise distance was lower in related pairs than in unrelated pairs, it was counted as a positive hit for related, and vice versa. The total number of genomes that fell in favor of related and unrelated were then compared. Similar analyses were performed comparing sharing of 1kB windows in assembled genomes. A genome was assigned a positive hit for related if the number of related pairs sharing the same strain of that genome exceeded the number of unrelated pairs sharing the same strain, and vice versa. To avoid artefacts arising from the fact that the number of unrelated pairs often vastly exceeds the number of related pairs, we downsampled each of the sets of unrelated pairs 100 times, resulting in the p-value distributions observed in Supplementary Figure 4.

Networks considered were spousal relationships (spouses versus non-spouse), household relationships (same versus different household), mother-child relationships (mother-child versus non-mother child), any social network connection (any connection versus no connection), and village (same versus different village). To ensure fair comparisons in the case of spousal relationships, a set of non-spousal pairs was constructed by considering all pairs possible between males of one marriage with females of a different marriage. Similarly, in the case of household relationships, a set of different household pairs was constructed by considering all pairs possible between members of one household and members of another. In addition, comparisons were also made between randomized networks of related and unrelated pairs, in which the identity of the network’s nodes were shuffled but the connections preserved, thus preserving the structure of the network.

### Social network predictions

For each pair of individuals, we created feature vectors containing the mean pair-wise SNP distance for each genome, the relative abundance of that genome in each individual, the number of shared genomes using 1kB outlier regions, and True/False values for whether a given genome was considered to be the same strain in both individuals using the 1kB outlier regions. These features were then used to train Random Forest Classifiers (RFCs) to predict spousal and household connections, where class-balanced datasets were constructed by downsampling the number of unrelated pairs to equal the number of related pairs (spouse/non-spouse; same household/different household). In order to train the RFCs on different data than those used in the predictions, we performed a 3-fold split of the related pairs and trained on two thirds of the data while predicting on the remaining third. Predictions from the three separate test sets were combined. ROC curves were constructed from the average of ten sets of 3-fold cross-validation, and p-values computed for the resulting AUCs using a Mann-Whitney U statistic on the confusion matrices.

## Data availability

Additional information on the samples can be obtained from www.fijicomp.org. All samples may be downloaded from NCBI Short Read Archive under Bioproject PRJNA217052. Note that the name for sample SRS475548 in SRA was incorrectly entered. It should be the oral microbiome sample from M2.33, not W2.33. All accession numbers are listed in Supplementary Table 1. Sample collection was voluntary, therefore not all of the individuals have oral and gut microbiome samples associated with the surveys.

Code for the analyses in this paper start with an alignment table in the form of a Python dictionary containing individual core genes as its highest level keys, where for each core gene there is a M x N x 4 numpy array, for M subjects, N loci, and four different alleles (A, G, C and T). Code for filtering these alignment tables into SNP tables, Manhattan distance calculations, and scripts for identifying non-shared mobile genetic elements from 1kb regions are posted on Github at https://github.com/thomasgurry/fijiComp_transmission.

## Acknowledgements

This work was supported by funding from the Center for Microbiome Informatics and Therapeutics at the Massachusetts Institute for Technology. This work was supported by grants from the National Human Genome Research Institute (U54HG003067) to the Broad Institute, the Center for Environmental Health Sciences at MIT, the Center for Microbiome Informatics and Therapeutics at MIT, and the Fijian Ministry of Health. I.L.B. is a Sloan Foundation Research Fellow, a Packard Fellowship in Science and Engineering, and a Pew Foundation Biomedical Scholar.

## Author contributions

This study was conceived of by I.L.B. and E.J.A. The study was designed by I.L.B., A.P.J., S.P.J., and E.J.A. Raw data was collected by I.L.B. and W.N. Metagenomic assemblies and metrics were developed and assessed by I.L.B., T.G., S.Z., K.H., S.K.Y., T.P.S., D.G.and E.J.A. Final analyses were performed by I.L.B. and T.G. The paper was written by I.L.B., T.G., and E.J.A. The authors have no conflicts of interest to report.

